# One step towards bioremediating a heavily polluted river: Metagenomic insights on the function of microbial community

**DOI:** 10.1101/675876

**Authors:** Luz Breton-Deval, Ayixon Sanchez-Reyes, Alejandro Sánchez-Flores, Katy Juárez, Patricia Mussali-Galante

## Abstract

The objective of this study is to understand the functional potential of the microbial community related to bioremediation activity and its relationship with the pollution of each site to enhance the future design of more accurate bioremediation processes. Water samples were collected at four sampling sites along the Apatlaco River (S1-S4), and a whole metagenome shotgun sequencing was performed to know and understand the microbial community involved in bioremediation. Additionally, HMMER was used for searching sequence homologs related to PET and polystyrene biodegradation and metal transformation in Apatlaco River metagenomes. The Apatlaco River is characterized by the presence of a broad spectrum of microorganisms with the metabolic potential to carry out bioremediation activities. Every site along the Apatlaco River has a particular community to perform bioremediation activities. The first site S1 has *Thiomonas, Polaromonas, Pedobacter, and Myroides,* S2 has *Pedobacter, Myroides, Pseudomonas and Acinetobacter, S3, Thiomonas, Myroides, Pseudomonas, Acinetobacter and Aeromonas; S4, Thiomonas, Myroides and Pseudomonas, Thauera.*

Furthermore, every site is rich in specific enzymes such as S1 has dioxygenase and dehydrogenase, which can degrade Catechol, Biphenyl, Naphthalene, and Phthalate. While, S2 and S3 are rich in dioxygenase and decarboxylating dehydrogenases to degrade Toluene, Fluorobenzoate, Xylene, Phenylpropanoate, and Phenol. S3 also has monooxygenases which degrade Benzene, and all the earlier mentioned enzymes were also found at S4.

## 1. Introduction

Rivers are complex systems closely and continuously connected to terrestrial ecosystems. The connection creates dynamic riverine ecosystems that experience a high flux of nutrients (Y. W. Zhao et al., 2019). Typically, the amount of Carbon (C) that rivers transport to the ocean is a small fraction of the total amount of C that rivers receive from terrestrial ecosystems. Some of the remaining C is returned to the atmosphere in the form of CO_2_, while another part is stored in river sediment (Wang et al., 2016). Besides, some C is also incorporated into the trophic chain. However, at present, rivers transport unusually high amounts of C and other compounds as a result of human activities in addition to the normal flow of C from terrestrial ecosystems (Peters et al., 2019).

A polluted river is a complex ecosystem given the different chemical substances that flow into the waterway such as heavy metals, hydrocarbons, high amounts of organic matter as well as chlorinated, nitroaromatic and organophosphate compounds (Y. Zhao et al., 2019). The biogeochemistry of polluted rivers is unbalanced because the energy captured to support either biosynthesis or respiration is more than the system requires; additionally, toxic compounds modify the microbial community (Li et al., 2013).

As a result, it is necessary to address pollution from a practical and affordable approach; a way to achieve this is via bioremediation. Bioremediation is a process used to reduce environmental contaminants by employing enzymes, microorganism, plants, microbial metabolites, or any bioproduct (Arora and Bae, 2014; Suthersan, 1999). However, a lot of uncertainty continues to surround some bioremediation strategies due to the lack of understanding of the specific microbes that are vital to performing the transformation of pollutants resulting from unspecific strategies that employ consortia of bacteria to carry out bioremediation projects (Dvořák et al., 2017). Only 1% of microorganisms are cultivable, and the community consortia have more microbes than the number of microorganisms that can be measured with traditional techniques (Ghosh and Bhadury, 2018). In contrast, Next-Generation sequencing such as the Whole Metagenome Shotgun (WMS) approaches to discover the total genomic content and microbial functions of the sample, which permits an understanding of the biochemical interactions of the microbial population with the surrounding environment without the limitations of the culture-based techniques. Besides, a resolution is higher than the universal 16S marker gene profiles (González-Toril and Aguilera, 2018; Techtmann and Hazen, 2016).

Research into the microbial ecology of impacted sites, including the complete genome sequences of diverse microorganisms and additionally their functions and the factors that influence these functions, could be useful to determine the gene pool of enzymes necessary for the degradation of the pollutant (Liu et al., 2018; Megharaj et al., 2011). Furthermore, this knowledge could help to select the best approach to eliminate pollution at the site. In some cases, the polluted site may include species that tolerate the contaminant, although this species is not dominant due to the lack of an appropriate carbon source (Techtmann and Hazen, 2016). In other contexts, there may not be a microorganism naturally available with the metabolic capacity to process the pollutant however a more effective microorganism could be added to the polluted location (Uhrynowski et al., 2017). A clear understanding of these characteristics of the river assists in the development of an effective bioremediation strategy to reverse the impact of pollution.

The Apatlaco River is located in central México in the state of Morelos. It is 63km in length and provides water to 10 state districts and an average of 2 millions of people (IMTA, 2008). The river receives 321 wastewater discharges, of which 49% come from industrial activities, 42% from the domestic sector and just 9% from farming (Fig. 1). As a result of these discharges, the contamination of the river is heterogeneous (Moeller-Chávez et al., 2004). Therefore, it is necessary to find a feasible solution to treat the river that is an essential key in regional development.

**Figure 1.**
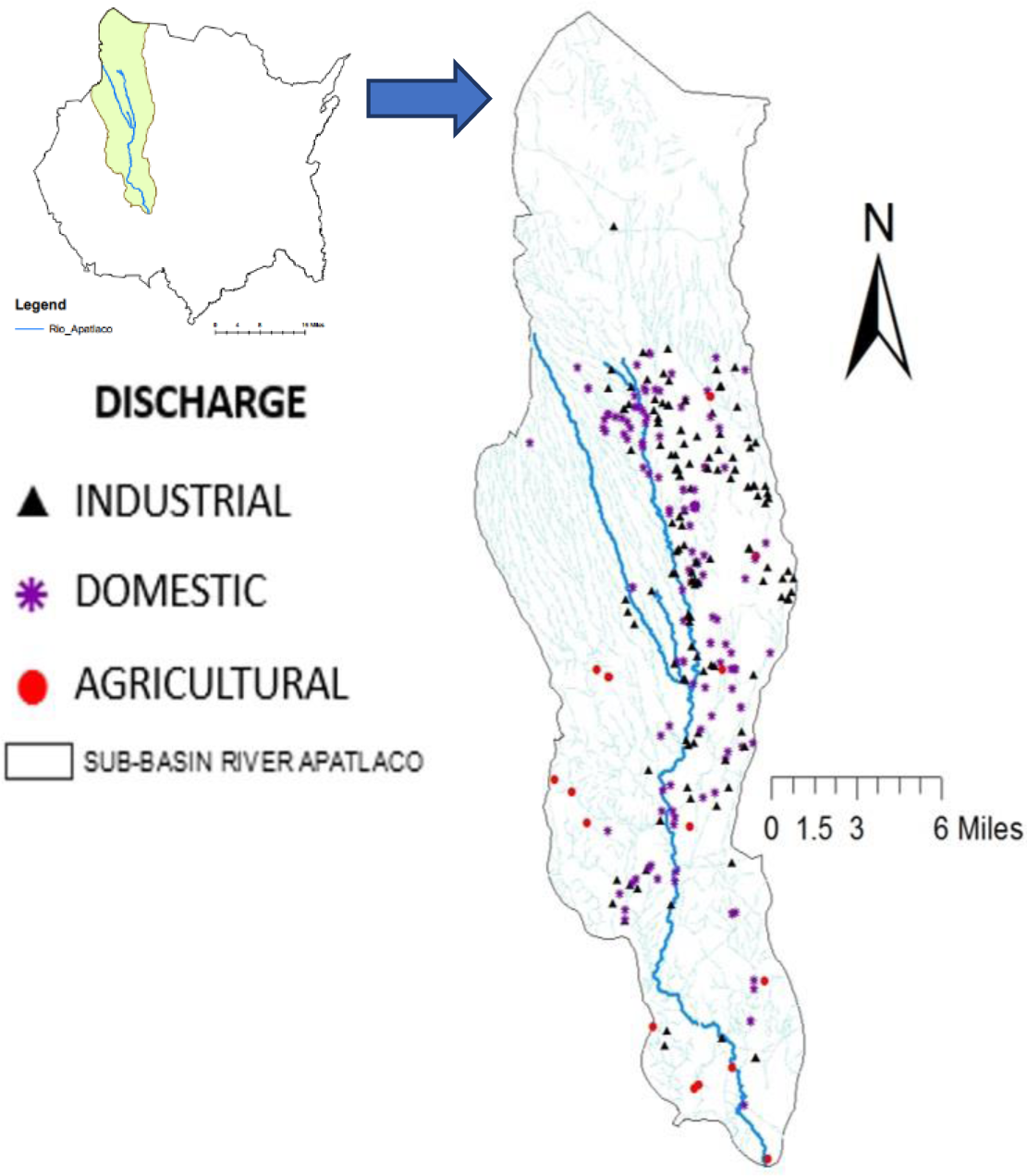
The Apatlaco River, where purple, black and red dots are the different discharges of domestic, industrial and agro origin respectively.

Therefore, the objective of this study is to understand the functional potential of the microbial community related with bioremediation activity and its relationship with the Apatlaco’s pollution to enhance the future design of more accurate bioremediation processes.

## Materials and Methods

### 1.1 Study site, sample collection, and chemical parameters measured

The water samples were collected at 4 sampling sites along the Apatlaco River (S1-S4), where S1 is clean, (−99.26872, 18.97372) and S2 (−99.2187, 18.83), S3 (−99.23337, 18.78971) and S4 (−99.18278, 18.60914) have different level of pollution (Table 1, Fig. 1). At each point, 10 different water samples of 1 L were collected and then combined to form a single sample for molecular analyses. After received, the samples were kept in ice and transported to the laboratory for further processing.

**Table 1.**
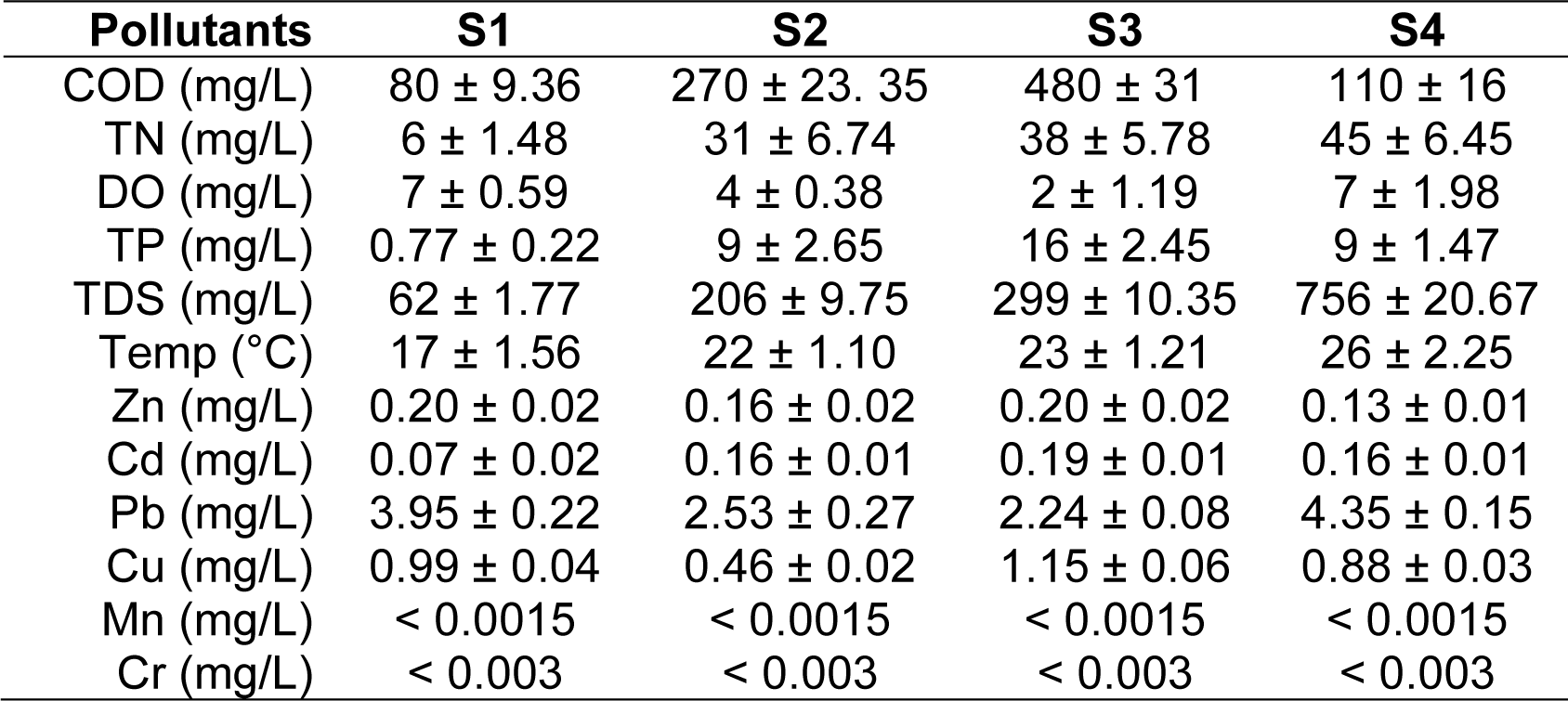
Measurements of physicochemical parameters along the Apatlaco River

HANNA multi parametric instrument DR900 was used to measure the following parameters: dissolved oxygen (DO), Total nitrogen (TN), total phosphorus (TP). The chemical oxygen demand (COD) was determined with the colorimetric method using HACH digester DRB200 and the DR900., and total dissolved solids (TDS) were measured according to Standard Methods.

Five water samples from Apatlaco River were analyzed for metal content by atomic absorption spectrophotometry (908 AA, GBC), with background correction. All samples were put to a final volume of 100ml. To ensure a satisfactory accuracy of the analysis, Standard Reference Material of National Institute and Technology and internal reference materials were used for precision, quality assurance, and control for selected metal measurements. All the material used was previously washed with HNO_3_ ultra-pure (J.T. Baker) for 24 hrs. For each measurement, the average values of three replicates were recorded. Metal content is reported as mg/L. (Zn, Cd, Pb, Cu, Mn, Cr). Detection limits of the atomic absorption spectrophotometer are: 0.0005 mg/L for Zn, 0.01 mg/L for Pb, 0.0015 mg/L for Mn, 0.003 mg/L for Cr, 0.001 mg/L for Cu, 0.0004 mg/L for Cd and 0.005 mg/L for Fe.

### 1.2 DNA extraction and Sequencing

DNA was extracted from water samples using a DNeasy PowerWater Kit (QIAGEN, Hilden, Germany). For each sample, an Illumina library was prepared from total DNA using the TruSeq kit v2 (Illumina, Inc., San Diego, CA, USA) following the manufacturer’s specifications with an average fragment size of 500 bp. The sequencing was performed on the NextSeq500 (Illumina, Inc., San Diego, CA, USA) platform with a 150-cycle configuration, generating paired-end reads with a length of 75 bp.

### 1.3 Data Analysis

After performing a quality control analysis using the FASTQC program and low-quality sequences were removed using an in-house Perl script (Andrews, 2010), the taxonomic profiling was performed using the raw reads with the software MetaPhlan v2.0 (Truong et al., 2017) using the following parameters: --bt2_ps sensitive-local --min_alignment_len 95 --input_typefastq. For the metagenome assembly, gene prediction and annotation, the raw reads were assembled using Megahit v1.1.3, Metagenemark v3.36, and Trinotate v3.1.1, respectively. The parameters used for Megahit were: --k-min 25 --k-max 75 --k-step 10 -m 0.4 -- no-mercy. The parameters used in Metagenemark were: -a -d -f G -m MetaGeneMark_v1.mod. For Trinotate, default parameters were used. GhostKOALA was used for KEGG’s annotation and K number assignment of metagenomic sequences (Kanehisha et al., 2016).

### 1.4 Biodegradative activities prediction by hidden markov models (hmm) profiles

HMMER was used for searching sequence homologs related to PET and polystyrene biodegradation. Furthermore, sequences involved in metals attenuation were mined as well in Apatlaco River metagenomes (da Fonseca et al., 2016). Were selected the most representative sequences belonging to:(i) 11 few well-characterized PET hydrolases (Danso et al., 2018), (ii) two sequences of styrene monooxygenase (StyA) functionally characterized and (iii) for metals were selected the well know transporters of Cd and Pb. After, for the construction of multiple independent alignments, we use Muscle (Edgar, 2009). The alignments were manually cured, and hmm profiles were generated with the hmm build command. Subsequently, were search with these models into the metagenomes obtained from Apatlaco River S1, S2, S3, S4 containing 499,126 sequences. We selected significant query matches with E-vale minimum of 0.001. Finally, hits with better E-values in each metagenome were selected for Blast search against the non-redundant NCBI database to assign potential taxonomies.

## Results and Discussion

### 2.1 Functional potential of the microbial community related to bioremediation in the Apatlaco River

The Apatlaco River is characterized by the presence of a broad spectrum of microorganisms with the metabolic potential to carry out bioremediation activities (Breton-Deval et al. 2019). The genera presented in table 2 are microorganisms located along the river that have been tested experimentally in previous research for their ability to tolerate or degrade pollutants (Gallego et al., 2006; Hovasse et al., 2016; Mattes et al., 2008; Mekuto et al., 2018).

**Table 2.** Microorganism with bioremediation potential

| Microorganism | Location | Biologicalactivity | Compounds | Pathogen | Ref. |
| --- | --- | --- | --- | --- | --- |
| *Thiomonas* sp. | S1, S3, S4 | Biofilms - Reduction | As, S, | No | (Zheng et al., 2014), 6-8 |
| *Polaromonas* sp. | S1 | Oxidation, biofilms | Hg, As,alkanes, Pyrene | No | (Mattes et al., 2008; Osborne et al., 2010; Singh et al., 2006; Yagi et al., 2009) |
| *Pedobacter* sp. | S1, S2 | Reduction desasimilative | As(V) | opportunistic | (Chen et al., 2016) |
| *Myroides* sp. | S1, S2, S3, S4 |  | CN<sup>-</sup>, SCN<sup>-</sup>Organic matter | opportunistic | (Mekuto et al., 2017; Samal et al., 2017) |
| *Pseudomonas* sp. | S2, S3, S4 | Biofilm, Biosorption | Pb +2, Ni +2, Cu +2, Cr +3, NO<sub>3</sub>, detergents, dyes, pesticides, hydrocarbons | opportunistic | (Naz et al., 2016; Ramadass et al., 2018; Roy et al., 2018; Wasi et al., 2013) |
| *Acinetobacter* sp. | S2, S3 |  | Hydroxydioxane, Cr, Clothianidin, cyprodinil, | yes | (Chen et al., 2018; Fernández et al., 2018; Wang et al., 2019; Zhang et al., 2017) |
| *Aeromonas* sp. | S3 | Biofilms - Siderophores | As, Cu, Fe, Ni, Zn, Mn (II), dyes, PHAs, | yes | (Chen et al., 2013; Du et al., 2015; Kumar et al., 2016; Uhrynowski et al., 2017; Zhang et al., 2019) |
| *Thauera* sp. | S4 |  | Zn, Cd, Co, Cu, Ni, Pb, Cr, Hg, Se | No | (Fang et al., 2018) |

In S1 the genera with these abilities were *Thiomonas, Polaromonas, Pedobacter, and Myroides*, the first three genera are specialists in heavy metal accumulation, biosorption, and reduction and the last one inorganic material mineralization (Magic-Knezev et al., 2009; Margesin and Zhang, 2013; Mekuto et al., 2018; Zheng et al., 2014). The *Polaromonas* genus found in S1 is representative of clean water with low levels of COD and nitrogen, just like S1 is (Table 1). The other genera perhaps are present due to the selective pressure of the high level of Pb and Cu. S2 exhibits genera including *Pedobacter, Myroides, Pseudomonas and Acinetobacter;* of which the last two have the metabolic machinery to degrade different compounds such as detergents, dyes, pesticides, hydrocarbons, clothianidin and cyprodinil (Ganguli and Tripathi, 2002; Paisio et al., 2016; Wang et al., 2019). In S2, all physicochemical parameters increased and DO decrease as a consequence of the rise of COD, the entire ecosystem has changed, and the microbial communities have been drastically modified. At this point of the river, the most abundant microorganism is *Acinetobacter* whose presence may be a result of waste discharge and growth prompted by the physicochemical conditions of the stream. Genera from the two previous sample points, including *Thiomonas, Myroides, Pseudomonas, Acinetobacter and Aeromonas,* are also present at S3. This point shows an increase in almost all levels of physicochemical parameters, except in Pb and OD. Finally, S4 includes *Thiomonas, Myroides and Pseudomonas* genera similar to earlier points in the river. However, *Thauera* genera are unique to this part of the river and have the potential to tolerate many metals including Zn, Cd, Co, Cu, Ni, Pb, Cr, Hg and Se (Fang et al., 2018). S4 is the last point before the river ends and the physicochemical conditions are very similar to the second sample point (S2) except for the higher levels of TKN and TDS due to the accumulation of nitrates, phosphorus, sodium, magnesium, and metals discharged along the length of the river.

### 2.2 Energy flux and the bioremediation potential

Microbial transformations of contaminants take place as a consequence of the use of pollutants as a source of carbon or as an electron acceptor (Fig. 2). O_2_ acts as the terminal electron acceptor in the process of aerobic respiration while NO_3_^−^, CO_2_ or fumarate can be used as terminal electron acceptors for anaerobic respiration. Aerobic respiration is energetically more efficient than other electron acceptors. However, O_2_ is not always the most available electron acceptor at all parts of the rivers. Despite this, the results showed evidence of microorganisms, which can use another electron acceptor such as nitrate. Regarding the aerobic degradation of pollutants predicted by the KEGG ortholog groups (KOs), there are several enzymes involved in the biodegradation of aromatic compounds. The enzymes found at S1 were dioxygenase and dehydrogenase, which can degrade Catechol, Biphenyl, Naphthalene, and Phthalate. While S2 and S3 are both rich in dioxygenase and decarboxylating dehydrogenases involved in the degradation of Toluene, Fluorobenzoate, Xylene, Phenylpropanoate, and Phenol. S3 also has monooxygenases which degrade Benzene. All of the earlier mentioned enzymes were also found at S4. These compounds are used to make lubricants, drugs, dyes, pesticides, and rubbers. Given that an industrial park is located close to the river, the afore mentioned presence of enzymes suggests that the industrial park does not have the necessary water plant to correctly treat their effluents and perhaps this constant discharge have selected the microbial community.

**Figure 2.**
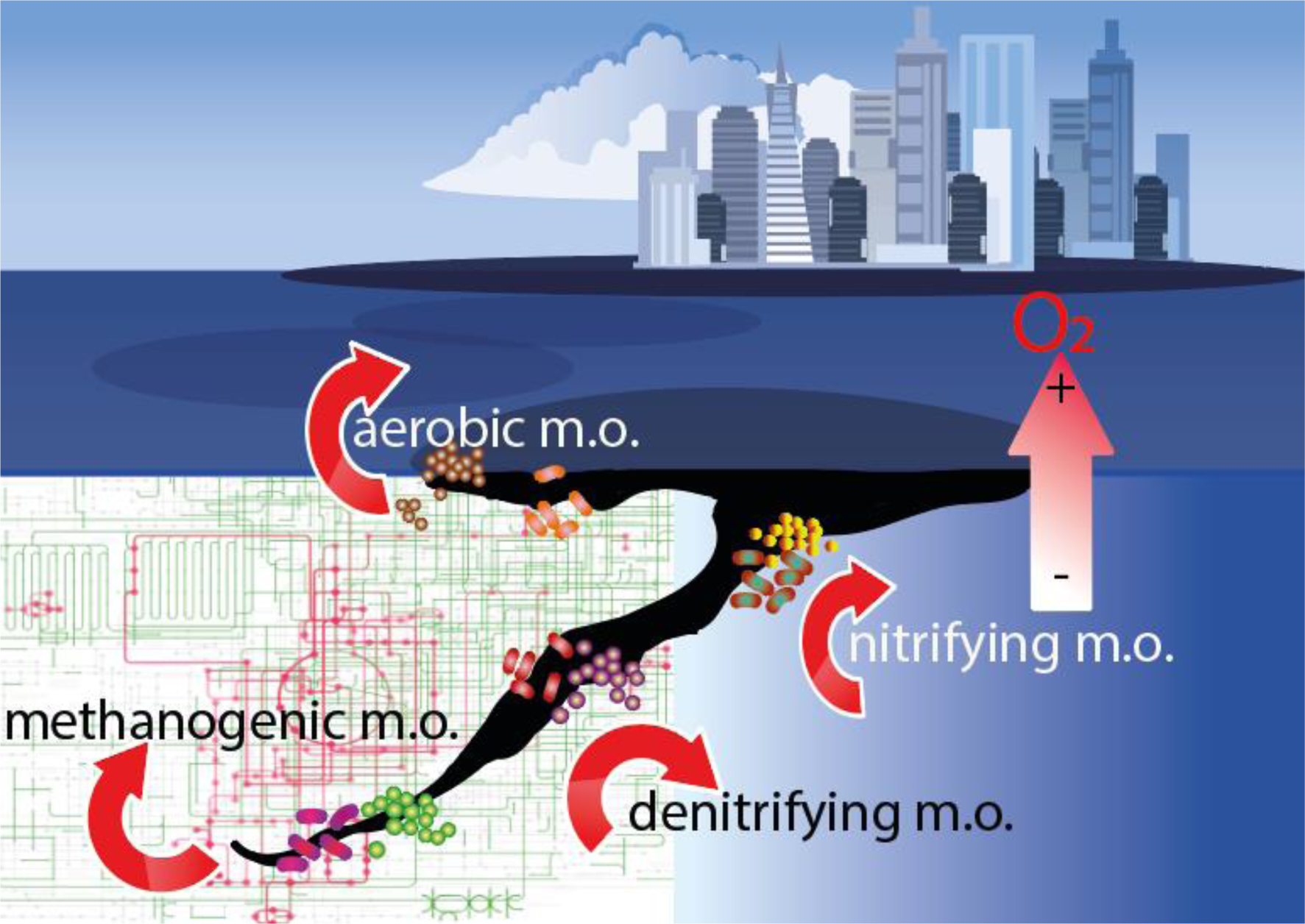
The dynamic microbial transformations of contaminants take place as a consequence of the use of pollutants as a source of carbon or as an electron acceptor

When O_2_ is scarce, nitrate is the next most energetically favorable electron acceptor which can reduce or denitrify bacteria; this last group is facultative anaerobes. The river is a dynamic system where several processes happen at the same time, some groups of bacteria use O_2_ while in oxygen deficient areas the nitrate reducing, or denitrifying bacteria use nitrogen. Figure 3 shows the logarithmic rise in the enzyme levels related to the increased level of TN. The enzymes involved in the nitrogen cycle were mainly oxidoreductases. Under the assumption of a different selective pressure gradient along the river, due to steady pollutant discharges, among S1-S4 sites we observe a differential distribution of the molecular functions assigned to various metabolic processes related to energy metabolism (Oxidative phosphorylation, Nitrogen metabolism, Methane metabolism; (Anexo A). Which suggests functional specialization in the communities that inhabit each point, and could determine the partition in a metabolic niche among different Apatlaco’s River prokaryotic communities that help to contend with anthropogenic contamination. These ultimate is maybe favorable in environments with dynamics resource availability.

**Figure 3.**
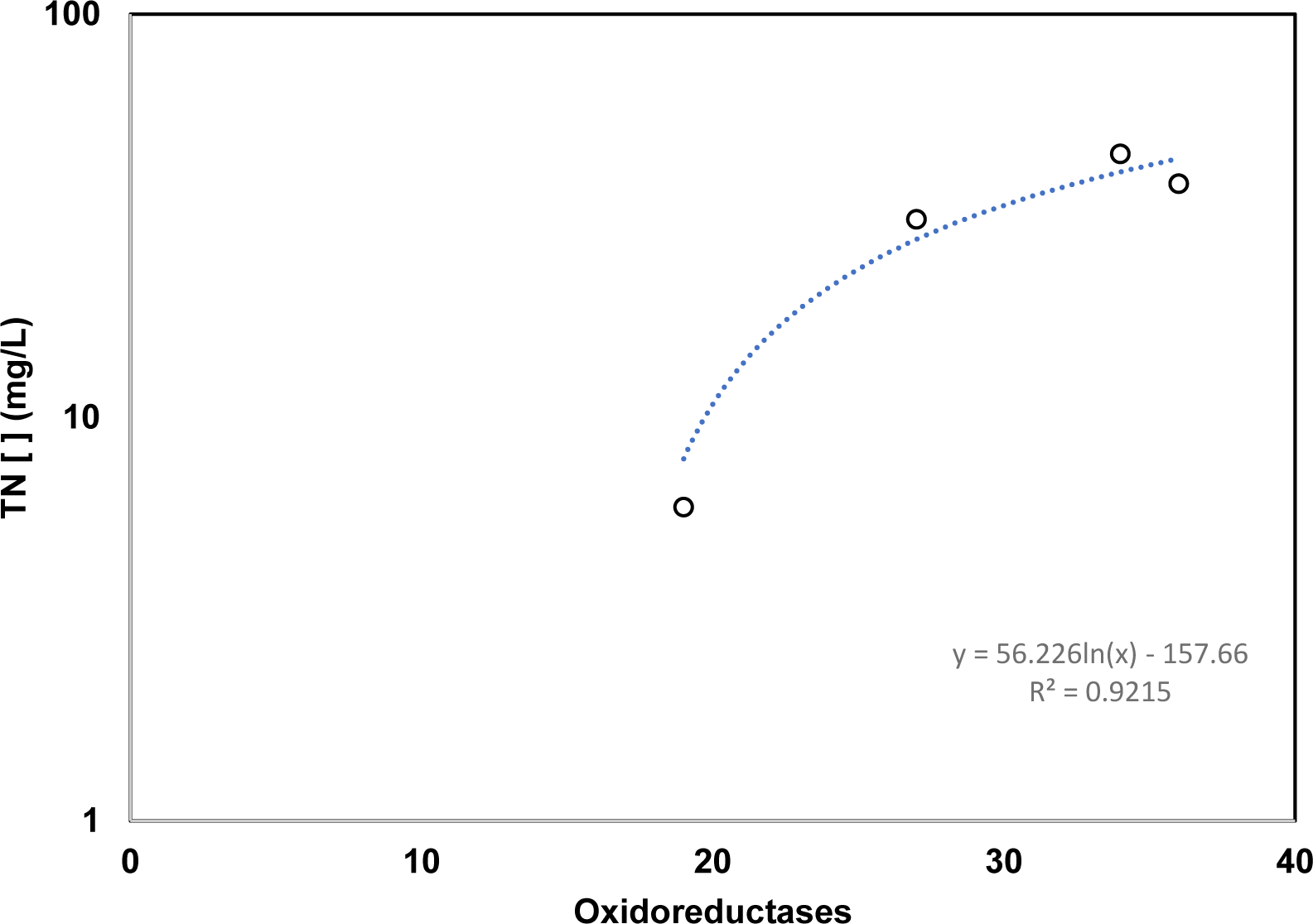
Logarithmic rise in the enzyme levels related to the increased level of Total nitrogen (TN).

Concerning the methanogenic pathway, the drivers are methanogens which produce methane as a byproduct and methanotrophs that consume that methane (Angelidaki W., 2004). The four sample sites have similar enzymes at particular points of the pathway such as the trimethylamine-N-oxide reductase (KO7811), a useful transmembrane electron carrier. However, there are specific enzymes at every point. S1 and S4 have genes for the hetero disulfide reductase (Hdr) required for the final reactions of the methane pathway. S1 also indicated the presence of methane monooxygenase (K16157), which is very useful for the degradation of alkanes, alkenes, ammonia, and aromatic compounds. At the same time, S2 has methanol dehydrogenases (K14028) to degrade ethanol, chloroethanol, and formaldehyde and S3 has a pyruvate ferredoxin oxidoreductase (K00169) useful for the degradation of nitrotoluene. This enzyme has been reported in *Methano bacterium thermo autotrophicum*, and in *Halobacterium halobium*, neither microorganism are not currently in that point however this enzyme is a homologous to a pyruvate ferredoxin oxidoreductase from *Thauera sp.* (Dörner and Boll, 2002) S4 has a methyl coenzyme M reductase (K00399) able to transfer sulfur groups. Concerning methanotrophs, all of the points showed trimethylamine-N-oxidoreductase (K07811-K07812) indicative of methane oxidation by methanotrophs.

### 2.3 Microbial genes and enzymes involved in the degradation of industrial pollutants: plastics and metals

The Apatlaco River is a basin mostly polluted with solid plastic residues such as PET derivatives and polystyrene foams among others. Every year the basin receives on average 5 tons of plastics (SEMARNAT, 2008) that remain without significant biological alteration. Although some databases include information about xenobiotic degradation pathways and the genes and enzymes involved in their degradation, some pollutants, such as PET and polystyrene, are missing. In order to find enzymes involved in PET and polystyrene degradation, Hidden Markov Model strategies were applied to search for potential homologous biodegradative sequences in several metagenomes obtained from The Apatlaco River (Anexo B). As expected, the PET hydrolase candidate’s genes belong to the superfamily of alpha/beta-hydrolases (pfamPF00561). Almost 50% of the sequences belong to the *Flavobacteriia* class. Interestingly, two sequences contain proteolytic domains suggesting similarities with the catalytic mechanisms of several peptidases. Eleven sequences appear to be classic esterase-lipase proteins, related to ester hydrolysis in a broad xenobiotic substrate family. Danso et al., 2018, retrieved two metagenomic PET hydrolase sequences from hmm probabilistic models that were functionally active in polycaprolactone and PET hydrolysis, supporting this approaching to undermine the metabolic potential in environmental microbiomes to degrade priority pollutants. Several studies suggest that the degradation of polystyrene by bacteria occurs by oxidizing individual units of styrene, through monooxygenase activities (Ho et al., 2018). We found six oxygenase (StyA) candidate’s genes (Anexo B). Homologous sequences in *Pseudomonas* and *Rhodococcusopacus* (Otto et al., 2004; Tischler et al., 2010) are responsible for activation in aerobic bacterial styrene degradation. We hypothesized that these metagenomic sequences could support the Apatlaco River’s microbial communities in the biodegrading of PET and polystyrene.

Other compounds that are missing in most databases include heavy metals, and this is because some metals such as Cu, Zn, and Fe are protein cofactors while other metals such as Cd, Ag and Pb do not play a role in bacterial metabolism (L. Wood et al., 2016). However, even at deficient concentrations, not essential metals or essential metals at high levels are toxic for microorganisms and it can be difficult to distinguish between the pathways involved in metabolism and the different strategies that microorganisms employ to deal with metals. Microorganisms can adsorb metals, change their speciation to less harmful compounds, and mineralize (Sharma and Pant, 2018). Biosorption is mostly carried out by proteins called metallothioneins while speciation is a common reduction mechanism; some examples are Cr^6+^ to Cr^3+^, AsO_4_^3−^ to AsO_3_^3−^, Hg^2+^ to Hg^0^ (Gutiérrez et al., 2018a).

The Aplataco’s water has some metals; however, only Pb and Cd levels were above the national standards (Table 1). Cd and Pb are metals that do not have any biological function and are very toxic because they may affect the renal, hematologic, and nervous systems. However, some reports of bacteria can tolerate 1350 mg/L of Cd and 1900 mg/L of Pb (Marzan et al., 2017). The genes involved in Cd resistance are RND exporters composed of the following proteins CzcA, CzcB, CzcZ, CzcN, CzcD, CzcR, CzcS which form transport systems that are able to export ions such as Zn2+, Co2+, and Cd2+ across the membranes. The application of Hidden Markov Model to find homologous sequences of these proteins allowed us the association with some microorganisms present in the river, with the ability to tolerate Cd (Anexo C). The number of species identified at each sample site is around five, except for S3 where 11 Cd tolerant species were found with a particular abundance of *Flavobacterium*. This same point has the highest level of Cd of the whole river (0.19 mg/L). It is possible that the elevated level of Cd stimulates a more significant presence of the microorganism. In the case of Pb, the genes involved in Pb resistance are metal-transporting ATPase for Cd, the ATPase for Cu+ CopA, the ATPase ZntA and the metallothionein protein smtA (Gutiérrez et al., 2018a). SmtA has been reported in *Synechococcus, Pseudomonas*, and *Yersinia pestis* (Gutiérrez et al., 2018b). While CopA and ZntA have been described in *E. coli* and *Enterococcus hirae* (Argudín et al., 2019). Our results identified microorganisms that haven’t been previously reported, such as *Limnohabitans, Cellvibrio, Polynucleobacter, or Azoarcus,* which can tolerate Pb (Anexo D).

## Conclusion

Our result allowed us to identify the microorganisms present along the Apatlaco River with the metabolic potential to carry out bioremediation activities of the following genera *Thiomonas, Polaromonas, Pedobacter, Myroides, Pseudomonas, Acinetobacter, Aeromonas and Thauera.* Furthermore, enzymes involved in the degradation of several priority pollutants were identified. The site S1 is rich in dioxygenase and dehydrogenase which can degrade Catechol, Biphenyl, Naphthalene and Phthalate. While, S2 and S3 are rich in dioxygenase and decarboxylating dehydrogenases to degrade Toluene, Fluorobenzoate, Xylene, Phenylpropanoate, and Phenol. S3 also has monooxygenases which degrade Benzene. All of the earlier mentioned enzymes were also found at S4. Although an oligotrophic stage prevails in the studied microbial communities of the Apatlaco’s River, the four points analyzed seem to show some specialization concerning energy metabolism, as well as the potential to obtain new biocatalysts for biodegradation of emerging pollutants such as plastic wastes. Finally, the bioinformatic analysis allowed us to identify microorganisms that haven’t been previously reported as a Pb tolerant (*Limnohabitans, Cellvibrio, Polynucleobacter, or Azoarcus)* for further research.

## Supporting information

Anexo

## Acknowledgments

The authors thank Instituto de Biotecnologia, UNAM for financial support to this research (P-9850). Also, LB-D thank to Consejo Nacional de Ciencia y Tecnología (CONACYT) and their program CATEDRAS for support the Project 285. We also like to thank the Unidad Universitaria de Secuenciación Masiva y Bioinformática (UUSMB) of the Instituto de Biotecnología, UNAM, for advice on DNA sequencing and Maria A. Millan for their help with the supplementary data. The map used in this paper was made by Karina E Rios-Ramos with data shared by the LISIG of the CIByC-UAEM.

