## Supplementary material for "One step towards bioremediating a heavily polluted river: Metagenomic insights on the function of microbial community": Anexo

**Anexo A**

**Table** Protein coding sequences differential abundance for Energy Metabolism according to KEGG annotation in S1, S2, S3, S4 microbiomes.

|  | **S1** | **S2** | **S3** | **S4** |
| --- | --- | --- | --- | --- |
| **Energy Metabolism** |  |  |  |  |
| Oxidative phosphorylation | **612**  (625.81)  [0.30] | **759**  (765.91)  [0.06] | **991**  (1050.37)  [3.36] | **770**  (689.91)  [9.30] |
| Nitrogen metabolism | **228**  (242.97)  [0.92] | **272**  (297.37)  [2.16] | **421**  (407.81)  [0.43] | **295**  (267.86)  [2.75] |
| Methane | **377**  (342.67)  [3.44] | **445**  (419.39)  [1.56] | **598**  (575.15)  [0.91] | **295**  (377.78)  [18.14] |
| Sulfur | **257**  (262.55)  [0.12] | **328**  (321.33)  [0.14] | **464**  (440.67)  [1.23] | **265**  (289.45)  [2.06] |

The contingency table provides the following information: the observed cell totals in black, (the expected cell totals) and [the chi-square statistic for each cell].

Chi-square statistic: 46.885.

p-value: < 0.00001.

**Anexo B**

**Table** Putative genes and enzymes involved in the degradation of PET and Polystyrenein Apatlaco´s river metagenome

| Metabolism | Degradation Pathways | Target Compound | Enzymes | Taxonomic assignation |
| --- | --- | --- | --- | --- |
| Plastic Metabolism | S1_gene_id_24935 | PET | S9 family peptidase | *Flavobacteriia bacterium* |
|  | S1_gene_id_74228 |  |  |  |
|  | S3_gene_id_78548 |  | OsmC family protein  Esterase | *Proteobacteria* |
|  | S1_gene_id_32163 |  | Alpha/beta hydrolase | *Bordetella flabilis* |
|  | S2_gene_id_115890 |  | dipeptidyl aminopeptidase | *Sphingobacteriales bacterium* |
|  | S4_gene_id_31632 |  | alpha/beta hydrolase | *Bacteroidia bacterium* |
|  | S2_gene_id_1309 |  |  |  |
|  | S4_gene_id_73202 |  |  |  |
|  | S3_gene_id_13623 |  |  |  |
|  | S3_gene_id_136251 |  | alpha/beta hydrolase | *Lutibacter*sp. |
|  | S1_gene_id_26234 |  | alpha/beta hydrolase | *Flavobacterium aquicola* |
|  | S2_gene_id_104464 |  | alpha/beta hydrolase | *Flavobacterium aquicola* |
|  | S3_gene_id_119837 |  | alpha/beta hydrolase | *Flavobacterium aquicola* |
|  | S1_sty_gene_id_71192 | Polystyrene | [kynurenine 3-monooxygenase](https://blast.ncbi.nlm.nih.gov/Blast.cgi#alnHdr_1085064631) | *Flavobacteria bacterium* |
|  | S1_sty_gene_id_12881 |  | FAD-dependent oxidoreductase | *Chitinophagaceae bacterium* |
|  | S2_sty_gene_id_93369 |  | Ferredoxin | *Aurantimicrobium*sp. |
|  | S2_sty_gene_id_14203 |  | FAD-dependent monooxygenase | *Flavobacteriia bacterium* |
|  | S2_sty_gene_gene_id_50356 |  | Ubiquinone biosynthesis protein UbiH | *Coxiellaceae bacterium* |
|  | S4_sty_gene_gene_id_61792 |  | Ferredoxin | *Aurantimicrobium* sp. |

**Anexo C**

**Table** Annotation of genes and microorganisms involved in tolerance to Cd

| **Gene** | **Site** | **Metal** | **Microorganism** | **KO** |
| --- | --- | --- | --- | --- |
| gene_id_11823 | S1 | Cd | *Flavobacteriumbranchiophilum* | K15726 |
| gene_id_14506 | S1 | Cd | *Methyloteneraversatilis* | K15726 |
| gene_id_29710 | S1 | Cd | *Methyloteneraversatilis* | K15726 |
| gene_id_41966 | S1 | Cd | *Emticiciaoligotrophica* | K15726 |
| gene_id_43320 | S1 | Cd | *Azospirillumsp. B510* | K01649 |
| gene_id_5825 | S1 | Cd | *Dechloromonasaromatica* | K15726 |
| gene_id_64667 | S1 | Cd | *Dechloromonasaromatica* | K15726 |
| gene_id_64869 | S1 | Cd | *Sphingopyxissp. QXT-31* | K15726 |
| gene_id_73324 | S1 | Cd | *Flavobacteriumbranchiophilum* | K15726 |
| gene_id_125204 | S2 | Cd | *Methyloteneramobilis* | K15726 |
| gene_id_19065 | S2 | Cd | *Chryseobacteriumtaklimakanense* | K15726 |
| gene_id_28653 | S2 | Cd | *Polynucleobacterduraquae* | K15726 |
| gene_id_58976 | S2 | Cd | *Cytophagahutchinsonii* | K15726 |
| gene_id_65914 | S2 | Cd | *Acinetobacterschindleri* | K15726 |
| gene_id_74637 | S2 | Cd | *Myroidesodoratimimus* | K15726 |
| gene_id_99674 | S2 | Cd | *Acinetobacterschindleri* |  |
| gene_id_14213 | S3 | Cd | *Acidovoraxcitrulli* | K15726 |
| gene_id_16557 | S3 | Cd | *Flavobacteriumanhuiense* | K15726 |
| gene_id_26792 | S3 | Cd | *Flavobacteriumcommune* | K15726 |
| gene_id_28643 | S3 | Cd | *Flavobacteriumcolumnare* | K15726 |
| gene_id_36296 | S3 | Cd | *Chryseobacteriumtaklimakanense* | K15726 |
| gene_id_3954 | S3 | Cd | *Azospiraoryzae* | K15726 |
| gene_id_45494 | S3 | Cd | *Polynucleobacterduraquae* | K15726 |
| gene_id_47077 | S3 | Cd | *Myroidessp. A21* | K15726 |
| gene_id_62056 | S3 | Cd | *Azospiraoryzae* | K15726 |
| gene_id_66897 | S3 | Cd | *Flavobacteriumcommune* | K15726 |
| gene_id_97580 | S3 | Cd | *Sulfurimonasautotrophica* | K15726 |
| gene_id_24337 | S4 | Cd | *Methyloteneramobilis* | K15726 |
| gene_id_61640 | S4 | Cd | *Runellaslithyformis* | K15726 |
| gene_id_64191 | S4 | Cd | *Polynucleobacterduraquae* | K15726 |
| gene_id_66902 | S4 | Cd | *Acidovoraxcitrulli* | K15726 |
| gene_id_73522 | S4 | Cd | *Shewanellasp. ANA-3* | K15726 |

**Anexo D**

**Table** Annotation of genes and microorganisms involved in tolerance to Pb

| **Gene** | **Site** | **Metal** | **Microorganism** | **KO** |
| --- | --- | --- | --- | --- |
| gene_id_3337 | S1 | Pb | *Flavobacterium johnsoniae UW101* | K01533 |
| gene_id_7468 | S1 | Pb | *Limnohabitans sp. 63ED37-2* | K17686 |
| gene_id_29294 | S1, S2 | Pb | *Limnohabitans sp. 103DPR2* | K17686 |
| gene_id_31926 | S1, S2 | Pb | *Limnohabitans sp. 103DPR2* | K17686 |
| gene_id_142524 | S2 | Pb | *Azoarcusolearius* | K17686 |
| gene_id_151010 | S2 | Pb | *Polynucleobacterasymbioticus* | K17686 |
| gene_id_28009 | S2 | Pb | *Acinetobacter johnsonii* | K17686 |
| gene_id_33717 | S2 | Pb | *Acinetobacter schindleri* | K17686 |
| gene_id_6128 | S2 | Pb | *Azoarcusolearius* | K17686 |
| gene_id_46638 | S3 | Pb | *Acidovorax sp. 1608163* | K17686 |
| gene_id_6116 | S3 | Pb | *Alicycliphilusdenitrificans BC* | K17686 |
| gene_id_91096 | S3 | Pb | *Alicycliphilusdenitrificans BC* | K01534 |
| gene_id_97562 | S3 | Pb | *Chryseobacteriumtaklimakanense* | K01533 |
| gene_id_99655 | S3 | Pb | *Acinetobacterjohnsonii* | K17686 |
| gene_id_12205 | S4 | Pb | *Cellvibriosp. PSBB006* | K17686 |
| gene_id_12330 | S4 | Pb | *Polynucleobacterasymbioticus* | K17686 |
| gene_id_14276 | S4 | Pb | *Limnohabitanssp. 63ED37-2* | K17686 |
| gene_id_47807 | S4 | Pb | *Beta proteobacterium CB* | K17686 |
| gene_id_66900 | S4 | Pb | *Limnohabitanssp. 63ED37-2* | K17686 |
| gene_id_73171 | S4 | Pb | *Azoarcusolearius* | K17686 |
| gene_id_80421 | S4 | Pb | *Acinetobacterjohnsonii* | K17686 |
